# The length constant, time constant and velocity of a propagating action potential

**DOI:** 10.64898/2026.06.05.728191

**Authors:** James A. Fraser, Eugenia Lopez-Belmonte Deza

## Abstract

Length and time constants are foundational to the study of conduction in neurons but are defined only for passive membranes. Here we define length and time constants for uniformly propagating action potentials. The derivation exploits transmembrane current transition (TCT) instants during action potential conduction when the net transmembrane ionic current is zero, but axial current remains non-zero. At these instants, we define a rate coefficient, *κ*, and from it define 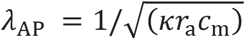 and *τ*_*AP*_ = 1/*κ*. We show that action potential propagation velocity is exactly *λ*_*AP*_/*τ*_*AP*_. We establish that *λ*_*AP*_ equals the ratio of axial to local capacitive current throughout the waveform, sets the exponential spatial weighting of ionic current, and is the local length scale of the *dV*/*dt* field at a TCT. It also approximates the length scale of the leading edge to within a fractional error of *τ*_*foot*_/*τ*_*m*_ under stated conditions. These identities are independent of channel gating formalism. We relate the new constants exactly to the passive DC, AC and AP foot constants and demonstrate the membrane and channel properties that determine *κ*. Finally, we suggest how *κ* may in principle be determined from experimental voltage waveforms.

## Introduction

Cable theory owes much of its enduring usefulness to two constants. In a passive cable, the length constant 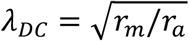 quantifies the distance over which the amplitude of a maintained voltage decays, while the time constant *τ*_*m*_ = *r*_*m*_*c*_*m*_ gives the rate at which a spatially uniform membrane charges [1]. Both describe the resting electrophysiological properties of a nerve axon and have been measured routinely since the 1940s [2]. A part of their value lies in their intuitiveness: as analytic solutions they allow the direction and approximate magnitude of the influence of their terms to be understood without formal calculation.

Analogous length and time constants for the propagating action potential (AP) have not been defined. This may reflect the fact that the amplitude of a propagating AP does not decay with distance, such that it is not immediately obvious what an AP length constant might quantify. Instead, the quantitative relationships between conduction velocity, the AP waveform and physical cable properties are described by the cable equation, into which Hodgkin and Huxley incorporated the non-linear dynamics of active ionic currents [3]. Rall and others subsequently extended cable theory to more complex geometries [4–6].

The active cable equation (derived in the Methods) quantifies the physical determinants of conduction velocity, *θ*, in uniform cables as:

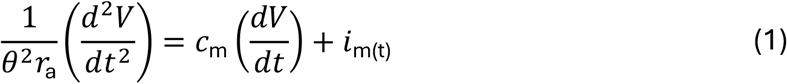

This identifies *r*_*a*_ (the axial resistance, *Ω* m^−1^), *c*_*m*_ (the membrane capacitance, F m^−1^) and *i*_*m*_ (the net transmembrane ionic current, A m^−1^) as the key determinants of conduction velocity, *θ* (m s^−1^), through their influence on the first and second differentials of membrane potential with respect to time. Note that here and throughout, transmembrane currents use the convention that a positive current denotes an outward electrical current, i.e. an outward flux of a positive ion or an inward flux of a negative ion.

The active cable equation has resisted expression in generalisable analytic form and consequently does not lend itself to the intuitive reasoning that the passive constants permit [7]. Because propagation depends on sharply voltage- and time-dependent conductances, it must generally be solved numerically [8], from the first digital solutions of the Hodgkin–Huxley system [9] to modern simulation environments such as NEURON [10,11]. Although analytical treatments have made substantial progress, they generally require simplified descriptions of the ionic currents [12–14].

In the absence of exact length and time constants for the active wavefront, two substitutes are in common use, each of which raises questions that the present work addresses. The first is to treat the AP foot as a single experimentally measurable exponential with exponent 1/*τ*_*foot*_ [1,3]. Its exact relationship to the rate of the subsequent active wavefront has not been established. The second substitute is the AC length constant, which captures the dominant influence of capacitance on the spatial voltage spread of APs, but requires determination of the frequency components of the AP [15]. It has also been employed to determine appropriate spatial discretisation of cable models [11]. However, a length scale specific to a given AP waveform would be useful both conceptually and computationally.

A further limitation of the lack of analytic forms is that numerical models provide forward descriptions of excitation and propagation but are poorly suited to the inverse problem of determining cable constants from experimentally recorded waveforms. Several specific questions therefore remain open. Can a propagating action potential be described in terms of length and time constants and, if so, what do they measure? How do such constants relate to the DC and AC length constants and to the constants of the AP foot? How do these active constants relate to membrane and cable properties? And can any of these quantities be recovered from an experimentally recorded voltage trace?

The present work answers these questions. We define a rate coefficient, *κ* and from it define AP time and length constants, *τ*_*AP*_ and *λ*_*AP*_. We establish their precise electrophysiological meaning. We relate the new AP constants exactly to the DC, high-frequency AC and AP foot constants. We use the latter relationship to suggest how *κ* may be determined from a voltage trace. We use computer modelling to determine how *κ* depends on membrane and ion channel properties, and show that quantities derived from *κ* report the erosion of conduction reserve before propagation fails.

All new derivations follow from the cable equation and are therefore independent of the formalisms used to represent the voltage-gated ion channels. We confirm this using a published six-state Markov description of *Na*_V_1.6 [16] and across parameter sets spanning the full range that supports propagation. We also test the correspondence between the new cable relationships and simulations using an electrodiffusive charge-difference model [17– 19].

## Methods

### The Cable Equation

A derivation of the linear cable equation is presented here for convenience and to allow comparison of its underlying assumptions with those of the computational model employed in this study. We first derive the equation in a general form that does not assume an ohmic membrane conductance, then introduce passive membrane resistance, *r*_*m*_, to consider the case of a passive cable.

During a propagating action potential, an axial current (*i*_*a*_) flows in regions where voltage (*V*) varies with distance (*x*), and is defined as positive in the direction of increasing *x*. By Ohm’s law, this implies:

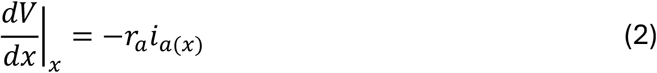

The axial resistance term, *r*_*a*_ (Ω m^-1^), strictly includes extracellular resistance, but as this is usually very much smaller than the true axial resistance we omit an explicit extracellular resistance term [1,4].

Resistive and capacitive transmembrane currents cause axial current to vary over distance, such that the change in *i*_*a*_ over distance must be equal and opposite to the local density of the total transmembrane current, *i*_*tot*_:

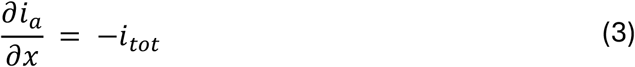

Differentiating equation (2) with respect to *x* gives:

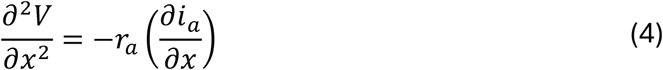

Then substituting equation (3) into (4):

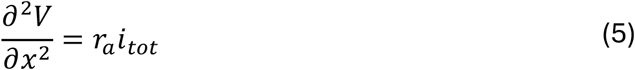

The total transmembrane current, *i*_*tot*_, has both capacitive and non-capacitive components. With outward current defined as positive:

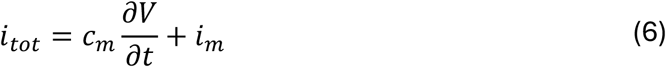

where *c*_*m*_ is membrane capacitance per unit length (F m^−1^) and *i*_*m*_ is the net non-capacitive transmembrane current per unit length. In the active cable, *i*_*m*_ is not assumed to obey an ohmic current-voltage relationship.

Combining equations (5) and (6) then gives the general active cable equation:

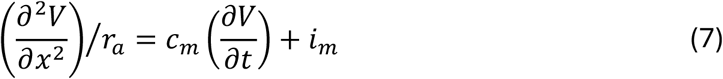

For an action potential waveform at constant velocity *θ*, the voltage at a point *x*_1_ at time *t*_1_ must be the same as the voltage at a point *x*_2_ at time *t*_2_, where *t*_2_ − *t*_1_ is the time taken for the wave to propagate from *x*_1_ to *x*_2_, so *x*_2_ = *x*_1_ + *θ*(*t*_2_ − *t*_1_). Thus, for the travelling waveform, *V*(*x, t*) = *F*(*z*):

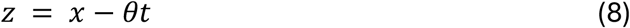

Using the coordinate in Equation (8), the chain rule applied to *V*(*x, t*) = *F*(*z*) gives:

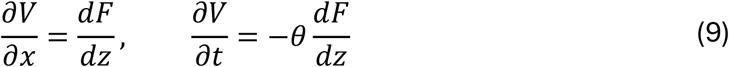

Thus:

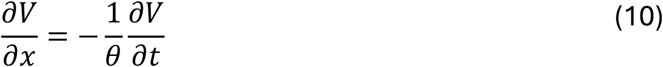

Differentiating once more gives the corresponding relationship between the second spatial and temporal derivatives,

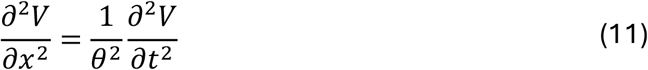

Substituting Equation (11) into (7) allows the active cable equation to be written in travelling-wave form as:

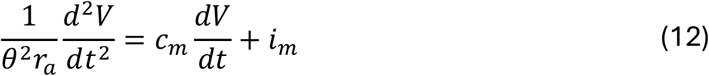

This is the form used in the present analysis. Note that this form of the cable equation allows non-linear variation in *i*_*m*_. In contrast, the familiar passive form of the cable equation requires the assumption that the membrane is passive and conductances are ohmic. Thus, where the membrane resistance *r*_*m*_ (Ω m) is the passive resistance around the resting potential such that *i*_*m*_ = (*V* − *V*_*rest*_)/*r*_*m*_, multiplying Equation (7) by *r*_*m*_ gives:

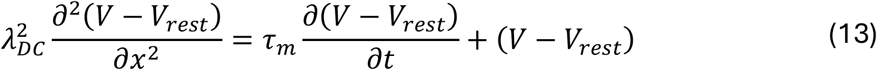

Applying the travelling-wave substitution (11) to equation (13) gives equation:

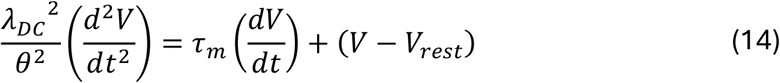

Where:

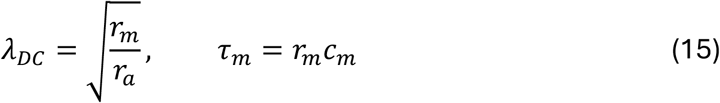

Thus, *λ*_*DC*_ and *τ*_*m*_ emerge only when the general active cable equation is specialised to an ohmic passive membrane and so, in contrast to Equation (12), do not describe the kinetics or scale of propagating AP waveforms.

### The charge-difference cable model

All simulations in this paper were conducted using the charge-difference cable model of Fraser & Huang [17,18], extended by [19]. This model was chosen because it does not share the simplifying assumptions inherent in the passive form of the cable equation, such as linear Ohmic current-voltage relationships and a lumped axial resistance term. Instead, the model employs first-principles physical formalisms. The model, previously employed to simulate skeletal muscle fibres, was reparametrised such that the base parameter set used in the present work approximates C-fibre vagal afferents [20], long unmyelinated axons. Definitions of the model parameters and variables and their default values are given in Table 1. For the baseline Hodgkin-Huxley simulations, voltage gated channel conductances were converted to permeabilities using the Goldman equation at the model’s steady state resting potential. The Markov validation instead retained conductance-based sodium and potassium currents, as described in the Supplementary Methods.

**Table 1.** baseline parameters.

| Symbol | Description | Value (units) |
| --- | --- | --- |
| $a$ | Fibre radius | 1.0 $\mu\text{m}$ |
| $c_m$ | Membrane capacitance per unit length | $6.28 \times 10^{-8} \text{ F m}^{-1}$ |
| $r_m$ | Membrane resistance = $R_m/(2\pi a)$ | $2.53 \times 10^5 \Omega \text{ m}$ |
| $r_a$ | Axial resistance per unit length | $2.28 \times 10^{11} \Omega \text{ m}^{-1}$ |
| $P_{\text{Na}}$ | Background $\text{Na}^+$ permeability | $2.16 \times 10^{-8} \text{ cm s}^{-1}$ |
| $P_{\text{K}}$ | Background $\text{K}^+$ permeability | $6.30 \times 10^{-7} \text{ cm s}^{-1}$ |
| $P_{\text{Cl}}$ | Background $\text{Cl}^-$ permeability | $1.00 \times 10^{-8} \text{ cm s}^{-1}$ |
| $P_{\text{H}_2\text{O}}$ | Membrane water permeability | $1.56 \times 10^{-3} \text{ cm s}^{-1}$ |
| $P_{\text{NaV(max)}}$ | Peak voltage-gated $\text{Na}^+$ channel permeability | $7.48 \times 10^{-4} \text{ cm s}^{-1}$ |
| $P_{\text{KV(max)}}$ | Peak voltage-gated $\text{K}^+$ channel permeability | $4.00 \times 10^{-4} \text{ cm s}^{-1}$ |
| $T$ | Absolute temperature | 300 K (27 °C) |
| $[\text{Na}^+]_e$ | Extracellular $\text{Na}^+$ concentration | 147 mM |
| $[\text{K}^+]_e$ | Extracellular $\text{K}^+$ concentration | 4 mM |
| $[\text{Cl}^-]_e$ | Extracellular $\text{Cl}^-$ concentration | 151 mM |
| $[\text{ATP}]_i$ | Intracellular ATP concentration | 6 mM |
| $[\text{ADP}]_i$ | Intracellular ADP concentration | 6 $\mu\text{M}$ |
| $[\text{Pi}]_i$ | Intracellular inorganic phosphate | 4.95 mM |
| $N$ | $\text{Na}^+/\text{K}^+$-ATPase density | $9 \times 10^{-12} \text{ cm}^{-2}$ |
| $\bar{A}_m$ | Opening rate factor, m gate | $273000 \text{ s}^{-1}$ |
| $\bar{B}_m$ | Closing rate factor, m gate | $3142 \text{ s}^{-1}$ |
| $\bar{V}_m$ | $\frac{1}{2}$ maximum voltage, m gate | -0.042 V |
| $V_{Am}$ | Opening voltage slope factor, m gate | 0.01 V |
| $V_{Bm}$ | Closing voltage slope factor, m gate | 0.018 V |
| $\bar{A}_h$ | Opening rate factor, h gate | $0.82 \text{ s}^{-1}$ |
| $\bar{B}_h$ | Closing rate factor, h gate | $6150 \text{ s}^{-1}$ |
| $\bar{V}_h$ | $\frac{1}{2}$ maximum voltage, h gate | -0.025 V |
| $V_{Ah}$ | Opening voltage slope factor, h gate | 0.011 V |
| $V_{Bh}$ | Closing voltage slope factor, h gate | 0.0136 V |
| $\bar{A}_n$ | Opening rate factor, n gate | $22900 \text{ s}^{-1}$ |
| $\bar{B}_n$ | Closing rate factor, n gate | $293.6 \text{ s}^{-1}$ |
| $\bar{V}_n$ | $\frac{1}{2}$ maximum voltage, n gate | -0.045 V |
| $V_{An}$ | Opening voltage slope factor, n gate | 0.007 V |
| $V_{Bn}$ | Closing voltage slope factor, n gate | 0.04 V |

### Key model formalisms

For the baseline Hodgkin–Huxley simulations, transmembrane currents for each ionic species, *s*, (Na^+^, K^+^ and Cl^-^) were calculated using the Goldman current equation [21]:

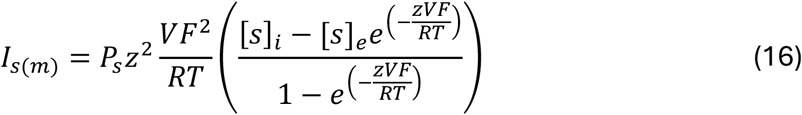

Axial conduction was calculated from the electrical current density form of the Nernst-Planck equation for each ion species, obtained by multiplying molar flux by *zF*:

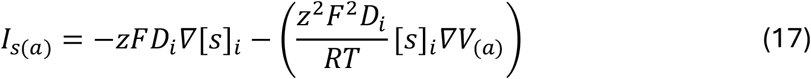

The membrane potential, *V*, was calculated in each compartment of volume *v* from the charge-difference equation [17,18]:

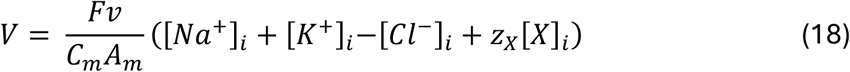

Other components of the baseline model, including Na^+^/K^+^-ATPase fluxes, ionic concentrations, volume changes and Hodgkin–Huxley voltage-gated channel kinetics, are as previously described [19], with the baseline parameters given in Table 1.

### Cable model discretisation

Unless stated otherwise, the baseline Hodgkin–Huxley simulations used a 4 mm cable. The 2 µm-diameter fibres in Figures 1 and 2 were represented by 444 longitudinal compartments (*Δx* = 9 µm); repeating these simulations with *Δx* = 5 µm produced no measurable change in conduction velocity or *κ*. Compartment length was reduced and compartment number increased for parameter sweeps as necessary to maintain *Δx* ≤ 0.1 *λ*_AP_. The Markov validation and spatial-wavefront simulations employed the figure-specific geometries and convergence checks described in the Supplementary Methods.

**Figure 1.**
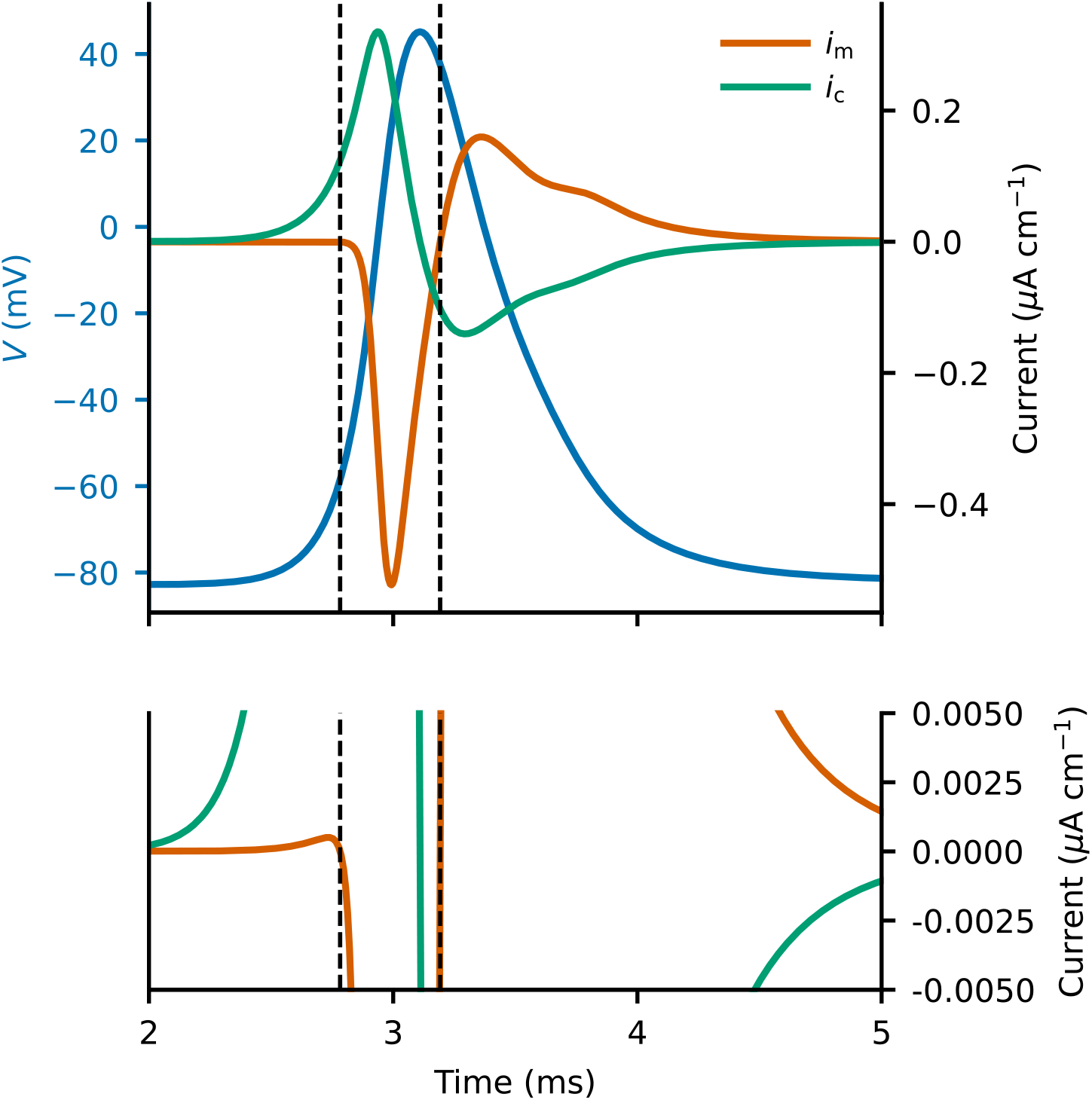
Transmembrane current transitions during a propagated action potential. Membrane potential V (blue), net transmembrane ionic current i_m_ (orange) and capacitive current i_c_ (green) recorded from the centre of a model 2 µm-diameter unmyelinated axon stimulated at one end at t = 0 ms. The lower panel shows the currents on an expanded scale, revealing the small outward i_m_ that precedes the much larger inward current. Vertical dashed lines mark TCT1, the outward-to-inward zero crossing of i_m_ during the upstroke, and TCT2, the inward-to-outward crossing during repolarisation. At both transitions i_m_ = 0 while the net axial and capacitive currents remain non-zero, such that dV/dt and d^2^V/dt^2^ are also non-zero.

**Figure 2.**
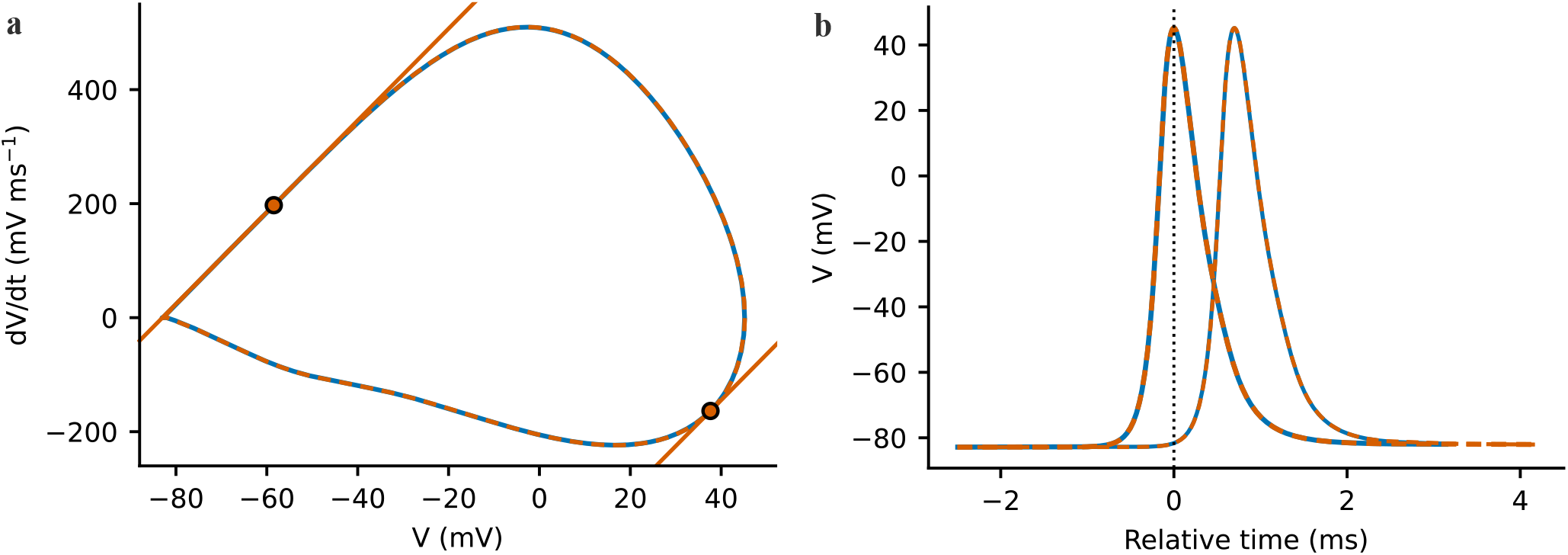
κ is invariant to fibre diameter when membrane kinetics and channel densities are unchanged. **(a)** Phase-plane trajectories 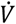 (V) for propagated action potentials in 2 µm-diameter (blue) and 1000 µm-diameter (orange dashed) model axons. Circles mark TCT1 and TCT2, and orange lines show the local tangents. All four tangents have slope κ = 8.045 ms^−1^. **(b)** Action potential waveforms at two sites in each axon, plotted relative to the peak time at the first site. Recording-site separations were 0.522 mm and 11.7 mm respectively, chosen to give comparable propagation delays. Measured conduction velocities were 0.748 and 16.7 m s^−1^ and agreed with 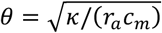.

### Stimulation and end effects

The baseline Hodgkin–Huxley cables used for Figures 1 and 2 were stimulated at one end with a 400 µs sodium-permeability pulse. Markov cables received a 50 µs permeability stimulus in the first two compartments. Recordings were analysed only within regions of uniform propagation. Uniformity was checked from local 0 mV-crossing velocity, peak voltage, action-potential duration, maximum upstroke rate and *κ* across compartments bracketing each reference site. The long Markov spatial simulation additionally established a continuous uniform region of 51.4 *λ*_AP_ encompassing the analysis region (Supplementary Methods).

### Calculation of *κ* from modelled action potentials

In simulations, TCT1 and TCT2 were identified from successive zero crossings of total transmembrane ionic current, from net outward to inward and from net inward to outward, respectively. *κ* was measured from the equivalent local phase-plane slopes 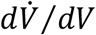 and 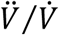. For the high-resolution Markov phase-plane and spatial reference, local derivatives at TCT1 were obtained from a quartic fit to *V(t)* over ±30 µs. For the Markov and Hodgkin–Huxley velocity comparisons, 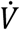 was first differentiated from the stored *V* samples, then an ordinary-least-squares line was fitted to 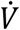 against *V* over the five saved waveform samples bracketing TCT1; this avoided the one-sided error in the online point estimate. Measured velocity was the reciprocal of the slope obtained by regressing linearly interpolated 0 mV-crossing times against distance across the reference compartment and its two neighbours on each side, with additional checks for uniformity at remote recording sites. The membrane resistance *r*_m_ was measured as the reciprocal centred slope of total membrane current with voltage for ±0.1 mV perturbations around rest, with ion concentrations and gating variables held fixed; *r*_a_ was obtained from the local axial voltage difference divided by the Nernst–Planck axial current per unit length. Note that this local determination of *κ* from current-defined TCTs is possible in simulations but does not represent a practical approach for determination of *κ* from experimental voltage traces, which is discussed in Section 5 of the Results.

The baseline parameter set is given in Table 1; parameters specific to the Markov validation are given in the Supplementary Methods.

## Results

### 1. Transmembrane current transition points and the rate coefficient, *κ*

#### 1.1 Identification and characterisation of transmembrane current transition points

Figure 1. displays a simulated action potential recorded from the central compartment of a model axon [17,19,20] stimulated at one end at time *t* = 0. The net transmembrane ionic current, *i*_*m*_, changes sign at two timepoints. We define these instants, at which *i*_*m*_ = 0 but the membrane potential is changing such that 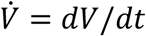 and 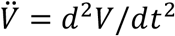 are non-zero, as transmembrane current transition (TCT) times.

The first TCT occurs when *i*_*m*_ turns from outward to inward. The initial small positive *i*_*m*_ develops as axial current from more proximal excited regions depolarises *V* locally, such that background outward currents (primarily *i*_*K*_, with smaller contributions from *i*_*Cl*_ and *i*_*Na*/*K*−*ATPase*_) initially exceed the inward Na^+^ current, *i*_*Na*_. However, *i*_*m*_ quickly turns negative as Na_*V*_ channels open and *i*_*Na*_ grows explosively. We define this first *i*_*m*_ = 0 crossing, at which 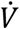 and 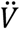 are non-zero, as TCT1. A second crossing, TCT2, occurs early during repolarisation, when *i*_*m*_ turns from negative to positive as outward currents again exceed inward currents, and at which 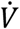 and 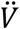 are again non-zero.

Complex waveforms and afterhyperpolarisations may contain additional crossings, but it follows from the description above that every uniformly propagating AP must contain TCT1 and TCT2 as defined.

#### 1.2 *κ* is the rate coefficient of the propagating wave

Rearranging Equation (12) gives:

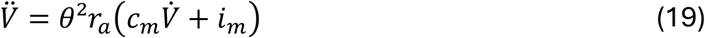

At a TCT, where *i*_*m*_ = 0 by definition, this simplifies to:

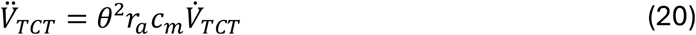

The ratio 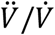 is defined at the TCTs because 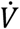 is non-zero there. The chain rule also identifies this ratio as the local slope of the phase-plane trajectory 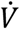 (*V*) and as the instantaneous logarithmic rate of 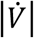. We therefore define:

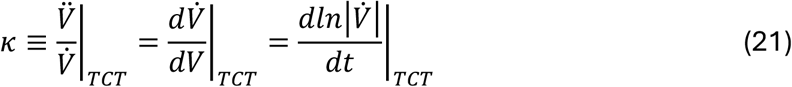

*κ* is therefore the rate coefficient of the propagating wave, with dimensions of time^−1^. Substituting Equation (20) into Equation (21) gives:

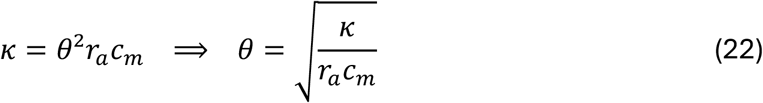

Equation (22) is exact for a uniform, shape-preserving travelling wave. By inspection, *κ* is therefore a single constant of the entire waveform, since *θ, r*_*a*_ and *c*_*m*_ are each single-valued for a given propagating AP. It follows that the phase-plane slope must be identical at every TCT of a given waveform. Importantly, it also follows that the same value of *κ* enters the quantitative description of every part of the AP waveform, as will be clarified below.

Figure 2 tests these predictions in model axons of 2 µm and 1000 µm diameter with identical membrane kinetics and channel densities. Despite the marked difference in diameter the two phase-plane trajectories overlap, and the tangent slopes at TCT1 and TCT2 were identical in both simulations, giving *κ* = 8.045 ms^−1^. Conduction velocities calculated from Equation (22) agreed with those measured from the propagation delay between compartments: 0.748 m s^−1^ for the 2 µm axon and 16.7 m s^−1^ for the 1000 µm axon.

#### 1.3 The cable relationships are independent of gating formalism

Equations (19)-(22) follow from the cable equation and the travelling-wave condition alone. These are independent of the mechanism generating *i*_*m*_ and so their derivations do not depend on the formalisms used to represent the voltage-gated ion channels. In order to confirm this central assertion, **Figure 3** tests the central *κ* – velocity relationship in an alternative channel model and with varied parameter sets. Thus, panel (a) shows propagating AP waveforms modelled using the published six-state Balbi Markov scheme for Na_V_1.6 and its associated Dodge–Cooley potassium formulation [16,22]. The waveforms contained both TCT1 and TCT2, and the same value of *κ* was recorded from the phase-plane tangents of each. An example phase plot is shown in panel (b). Na_v_ density and membrane capacitance were varied across the tested range of parameter combinations that produced successful AP propagation and a stable resting potential. For all such simulations, *κ* measured from the waveform predicted conduction velocity by Equation (22) over a >5-fold range. Similar measurements were made over Hodgkin–Huxley parameter sets spanning the full range that supported stable propagation (panel d). Over a >7-fold range of velocities, Equation (22) predicted the measured conduction velocity well.

**Figure 3.**
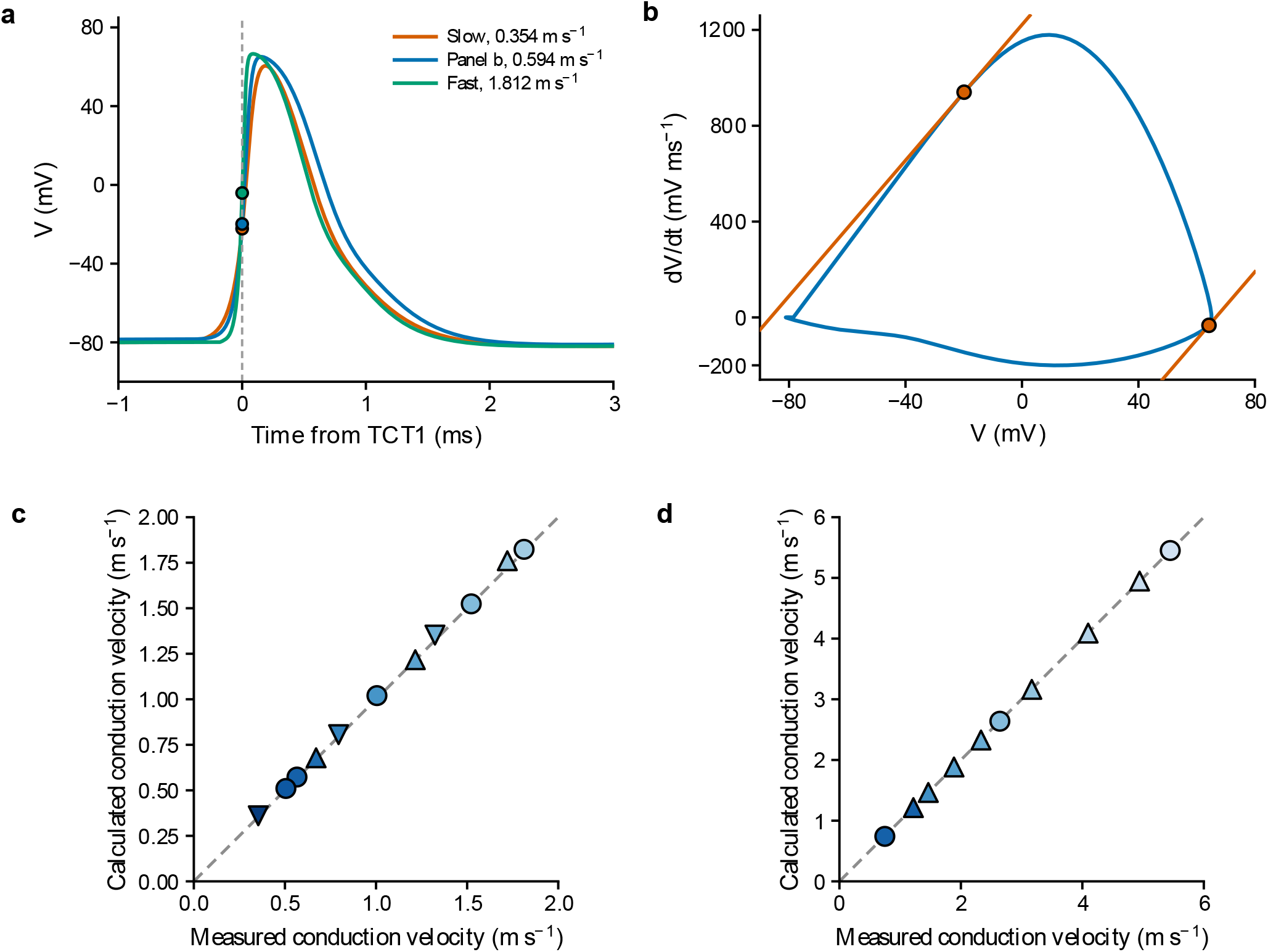
Velocity is calculable from κ independent of gating model. **(a)** Propagated action-potential waveforms for the condition used in panel b and the slowest and fastest uniformly conducting conditions in the validated Markov NaV-capacitance sweep. Time plotted relative to TCT1 for each waveform; circles mark each waveform’s TCT1. **(b)** Phase-plane trajectory for a propagated action potential generated using the six-state Balbi Markov scheme for NaV1.6. Orange lines are tangents at TCT1 and TCT2 and the measured slope at both is κ = 14.143 ms^-1^. **(c, d)** Conduction velocity calculated as 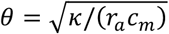 against velocity measured from interpolated 0 mV crossing times for 11 independently equilibrated Markov model parameter sets (c), and for ten Hodgkin-Huxley parameter sets (d). In both c and d, membrane capacitance increases with the intensity of blue shading; NaV density is indicated by downward triangles, circles and upward triangles for decreased, baseline and increased, respectively. The identity line is shown as a grey dashed line in both panels.

Note that although the cable relationships are independent of the ion channel formalism used and *κ* has a definition that is independent of gating formalisms, the numerical value of *κ* is a waveform measure, some determinants of which are explored in Section 4 below.

### 2. *κ* defines the AP time constant, length constant and conduction velocity

#### 2.1 The action potential time and length constants

The familiar passive constants describe voltage spread in the special case of unchanging ohmic conductances. The corresponding length and time scales of active propagation follow from *κ*. Because *κ* is the instantaneous logarithmic rate of 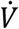 at a TCT, its reciprocal provides a natural definition of the AP time constant as:

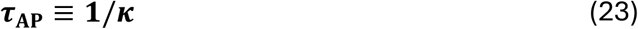

For the travelling coordinate *z* = *x* − *θt*, the temporal and spatial derivatives of any waveform quantity are related by ∂/ ∂*t* = −*θ* ∂/ ∂*x*. From Equation (22) the corresponding spatial constant is therefore:

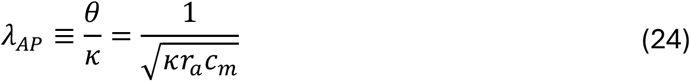

Writing the electrical diffusivity of the cable as *D* = 1/(*r*_*a*_*c*_*m*_) then gives

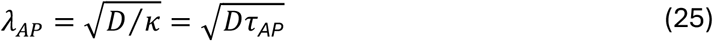

Conduction velocity and cable diffusivity are then exactly related to the AP constants:

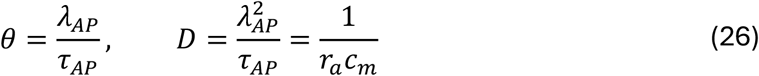

A uniformly propagating AP therefore travels one *λ*_*AP*_ in one *τ*_*AP*_. For the 2 µm-diameter model axon of Figures 1 and 2, *κ* = 8.045 ms^−1^ gives *τ*_*AP*_ = 0.1243 ms and *λ*_*AP*_ = 0.0932 mm, with *λ*_*AP*_/*τ*_*AP*_ = 0.750 m s^−1^, compared with the directly measured velocity of 0.748 m s^−1^.

It is important to define the physical meaning of *λ*_*AP*_, since the amplitude of a uniformly propagating AP does not decline with distance. The aspects of a propagating AP that may be quantified by *λ*_*AP*_ are established in the following three subsections.

#### 2.2 *λ*_*AP*_ is the ratio of axial to capacitive current throughout the waveform

The relationships in Section 2.1 can be expressed in terms of axial current by Ohm’s law to derive a first interpretation of *λ*_*AP*_. For a positive-going travelling wave, 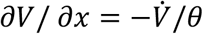, thus:

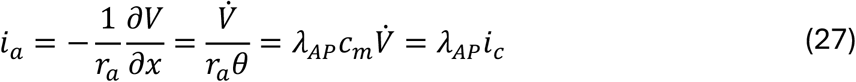

We recognise 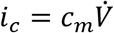 as the capacitive current per unit length. Thus:

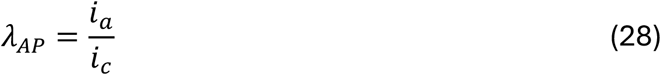

Thus *λ*_*AP*_ is the ratio of net axial current to local capacitive current per unit length wherever 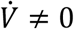. The dimensions confirm that this is a length: *i*_*a*_ is in A whereas *i*_*c*_ is in A m^-1^. This proportionality holds throughout the waveform.

This is the exact active counterpart of the corresponding relationship in a passive cable. For a passive steady exponential decay of voltage from its source, *i*_*a*_ = (*V* − *V*_∞_)/(*r*_*a*_*λ*_*DC*_) = *λ*_*DC*_*i*_*m*_. The AP constant replaces that steady-state leak term with a capacitance term to describe the travelling wave.

#### 2.3 *λ*_*AP*_ is the range over which transmembrane ionic current sets the local rate of change of membrane potential

Because *κ* is single-valued for a given propagated waveform, Equation (22) may be used to rewrite Equation (19) and the result written in terms of *κ* alone. This gives a relationship for the entire waveform, including but not limited to the TCTs:

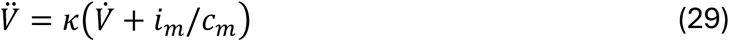

The homogeneous solution, obtained by setting *i*_*m*_ = 0, is 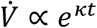. Thus *κ* is the instantaneous logarithmic rate of 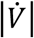 at a TCT, and elsewhere *i*_*m*_ drives its value away from this reference exponential. For a wave travelling in the positive *x* direction, ∂/ ∂*t* = −*θ* ∂/ ∂*x*. Substitution gives the exact spatial counterpart as a first-order relation for 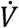:

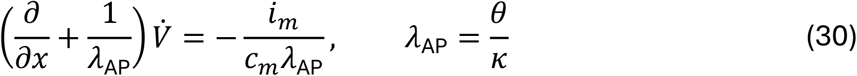

Multiplication by the integrating factor 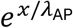, followed by integration from the fully repolarised membrane far behind the wave, where 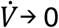 as *x* → −∞, gives:

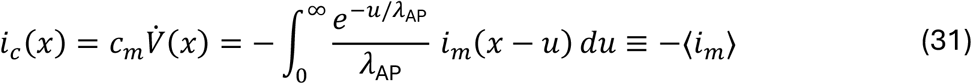

The kernel integrates to unity, so ⟨*i*_*m*_⟩ is an exponentially weighted average of the ionic current over the membrane behind the observation point. The local capacitive current is equal and opposite to this average. At TCT1, the local ionic current is zero, yet capacitive charging continues: the weighted ionic current behind that position is inward.

*λ*_AP_ is both the e-folding length and the mean spatial lag of the weighting: 63% of the weight lies within one *λ*_AP_ and 95% within three. This gives *λ*_AP_ an exact spatial meaning throughout the travelling wave, including regions where neither voltage nor its time derivative is exponential.

The comparison with *λ*_DC_ is instructive. In an infinite uniform passive cable, a maintained point current injected at *x* = *s* produces the steady voltage profile [1]:

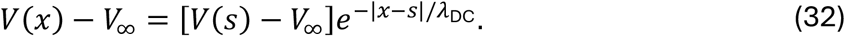

Here *λ*_DC_ is the e-folding length of a symmetric kernel, with current spreading in both directions from the source. Both constants therefore characterise exponential spatial weighting: *λ*_DC_ describes the passive current and voltage response to an applied current, while *λ*_AP_ describes the active ionic and capacitive currents within the travelling wave. This complements the current-ratio interpretations of Section 2.2: *λ*_AP_ = *i*_*a*_/*i*_*c*_ in the travelling wave, and *λ*_DC_ = |*i*_*a*_|/|*i*_*m*_| in either direction away from the source.

#### 2.4 *λ*_*AP*_ is the local length scale of the *dV*/*dt* field at a TCT

For any non-zero field *f* carried by a positive-going travelling wave, its signed local length is the reciprocal of its negative logarithmic spatial slope. Using the travelling-wave relationship (Equation (10)), with Equations (24) and (29) allows for an exact expression for the length of the 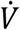 field, 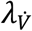:

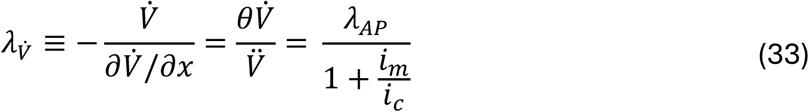

Equation (33) may be interpreted as quantifying the partition of current. Because *i*_*a*_ and *i*_*c*_ are both proportional to 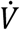, their local spatial scales are the same. At a TCT, where *i*_*m*_ is zero, the local spatial scale of both is exactly *λ*_*AP*_.

Other than at the TCTs, the discrepancy of 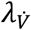 from *λ*_*AP*_ is determined by the ratio between *i*_*m*_ and *i*_*c*_. During the accelerating part of the AP upstroke, where 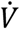 and 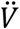 are positive, outward *i*_*m*_ shortens the local scale, whereas inward *i*_*m*_ increases it. At maximum 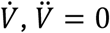 and the local scale diverges.

#### 2.5 The AP foot constants follow exactly from *κ*

Sections 2.2 to 2.4 establish the aspects of the waveform that *λ*_*AP*_ describes directly. We now show how it relates to the time and length scales of the AP foot, a part of the AP that has a long history of direct measurement [1]. Thus, the early leading edge of a propagating AP may be characterised by a time constant, *τ*_*foot*_, fitted to the exponential rise that precedes the main upstroke.

The AP foot may be defined as the period during the early leading edge in which the net transmembrane ionic current can be represented as *i*_m_ = (*V* − *V*_∞_)/*r*_m_, with *r*_m_ and *c*_m_ effectively constant. This is the classical ohmic-foot assumption [1], under which the relationships below are exact. Throughout, *τ*_m_ = *r*_m_*c*_m_ is the passive membrane time constant.

Let *V* − *V*_∞_ ∝ *e*^*αt*^ during this interval, such that *α* ≡ 1/*τ*_*foot*_. Substitution into Equation (29) gives:

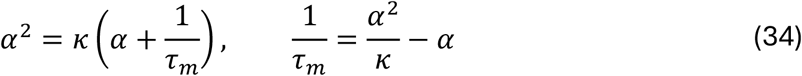

Since *τ*_*m*_ is positive, the foot exponent *α* always exceeds *κ. τ*_*foot*_ is correspondingly shorter than *τ*_*AP*_, and the two are related exactly by rearrangement of Equation (34):

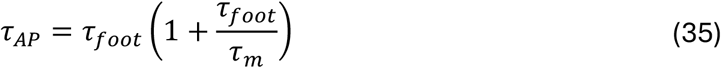

Defining the spatial constant of the foot as the travelling-wave counterpart of its time constant by *λ*_*foot*_ ≡ *θτ*_*foot*_ gives the corresponding relationship between lengths:

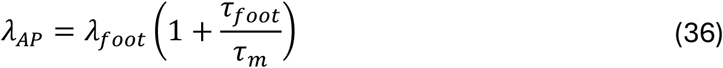

The foot constants are therefore shorter than their AP counterparts by the same fractional amount, *τ*_*foot*_/(*τ*_*m*_ + *τ*_*foot*_), which approaches *τ*_*foot*_/*τ*_*m*_ when *τ*_*foot*_ ≪ *τ*_*m*_. Equation (34) may be solved explicitly in either direction, giving forms that may be useful experimentally, as will be explored in Section 5:

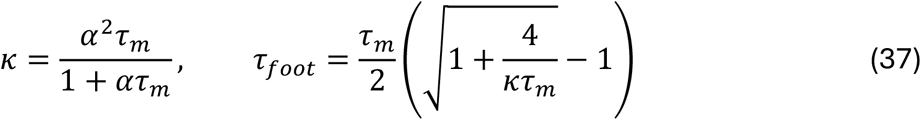

These relationships have a direct current-partition interpretation that connects them to Section 2.2. For the exponential foot, the fractions of converging axial current carried by capacitive charging and by the resting ionic conductance are, from Equations (34)-(36):

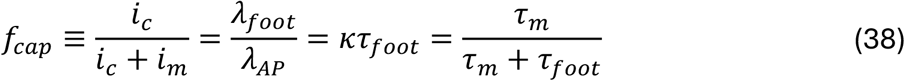

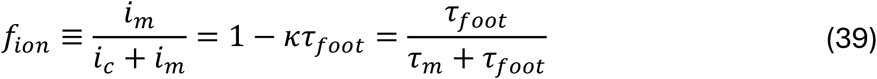

Under the ohmic-foot assumption, the converging axial current has only these two destinations, so *f*_*cap*_ + *f*_*ion*_ = 1. The single dimensionless product *κτ*_*foot*_ therefore states both what proportion of the foot current is capacitive, and how closely *λ*_*foot*_ approaches *λ*_*AP*_. Where the resting conductance is a small fraction of the total, *κτ*_*foot*_ approaches unity and the measured foot exponent closely approximates *κ*; where resting leak is more substantial, *λ*_*foot*_ becomes systematically shorter and the two diverge.

#### 2.6 Simulations of the exact relationships between *λ*_*AP*_ and the waveform

In Sections 2.2 to 2.5 we have shown the relationship between *λ*_*AP*_ and the axial current, capacitive currents, and the AP foot constants. In this section we show how closely each of these predictions are met in Hodgkin-Huxley and Markov model fibres.

**Figure 4**, panels **(a)** and **(b)** demonstrate a test of Equation (36), which quantifies the relationship between *λ*_*AP*_ and *λ*_*foot*_ from two measurable parameters, *τ*_*foot*_ and *τ*_*m*_. The two model fibres depicted have values of *τ*_*foot*_/*τ*_*m*_ that are some 20-fold different. Both panels show AP waveforms from compartments separated by 1 or 2 x *λ*_*AP*_ (left panels) and the corresponding spatial waveform (right panels) from every compartment of the plotted range at the TCT1 time, giving a spatial profile of *dV*/*dt* ahead of and behind TCT1.

**Figure 4.**
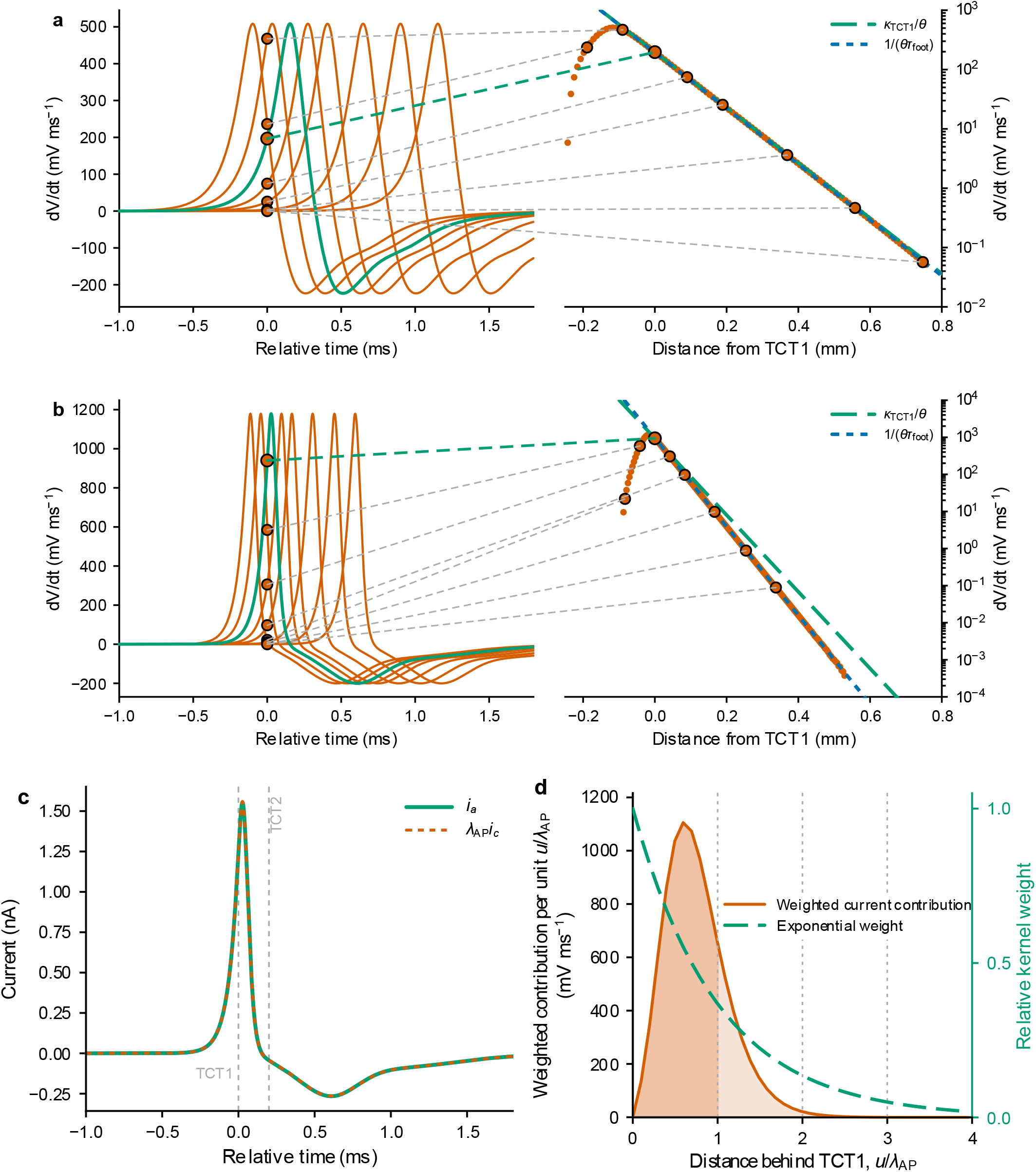
λ_AP_ predicts λ_foot_, axial current and the range of its influence. **(a, b)** dV/dt (left panels) and corresponding spatial waveform of positive dV/dt values plotted on a logarithmic scale (right panels) for (a) the Hodgkin–Huxley fibre of Figures 1 and 2 and (b) the Markov fibre of Figure 3. In the left panels, propagated AP waveforms are shown over time in selected model compartments at −2, −1, 0, 1, 2, 4, 6 and 8 λ_AP_ from the reference compartment (green). In the right panels, the spatial waveforms of these APs at the TCT1 time of the reference compartment are shown for all modelled compartments. Connectors indicate the relationship between the selected waveform points in the left and right panels, with the connector from TCT1 of the reference compartment highlighted in green. In the right panels, long green dashes are tangents at TCT1 and indicate the reference logarithmic decay rate κ/θ = 1/λ_AP_; short blue dashes show the foot decay rate 1/(θτ_foot_) = 1/λ_foot_. The foot decay rate exceeds κ/θ by 15.6% in the Markov example and by 0.76% in the HH example. Panel **(c)** shows the axial current trace of the reference compartment of panel b and compares this with the axial current calculated from i_a_ = λ_AP_c_m_dV/dt. Panel **(d):** Spatial reconstruction of local dV/dt at TCT1. At each distance u behind the observation point x_0_, the ionic current per unit cable length i_m_(x_0_ − u) was weighted and divided by c_m_. The orange curve shows 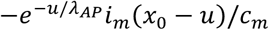, the contribution to local dV/dt per unit u/λ_AP_. Integration over the membrane behind the observation point gives 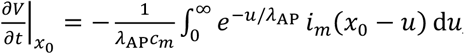, the shaded area. This gave dV/dt = 940.84 mV ms^-1^ compared to 940.11 mV ms^-1^ measured directly at x_0_. Darker shading denotes the contribution from within the first λ_AP_ behind TCT1 and paler shading the remainder. The green dashed curve shows the relative exponential weight; grey lines mark successive −λ_AP_ intervals.

**Figure 4a** simulates a fibre with the Hodgkin–Huxley parameters used in Figs 1 and 2. Its high passive membrane resistance and high excitability gives *τ*_*m*_ = 15.87 ms, *τ*_*foot*_/*τ*_*m*_ = 0.00778 such that *λ*_*AP*_ and *λ*_*foot*_ are predicted to differ by only 0.78%. The two spatial slopes are correspondingly indistinguishable across four decades of *dV*/*dt*, and the measured ratio between them was 1.0076, in agreement with the prediction to within 0.02%. This also confirms the prediction from Equation (36) that the distinction between the AP and foot length scales is negligible in fibres of high passive resistance.

**Figure 4b** depicts the Markov model fibre used in Figure 3. This fibre has much lower passive membrane resistance giving *τ*_*m*_ = 0.371 ms, measured *τ*_*foot*_ = 61.13 µs, thus *τ*_*foot*_/*τ*_*m*_ = 0.165. Equation (36) therefore predicts that *λ*_*AP*_ and *λ*_*foot*_ should differ by 16%. At TCT1 the local spatial decay rate of the 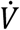 field was *κ*/*θ* = 1/*λ*_*AP*_, giving *λ*_*AP*_ = 41.97 µm, in agreement with Equation (33). In the waveform ahead of TCT1 the profile noticeably steepens, converging over approximately one *λ*_*AP*_ onto a straight far-foot asymptote of slope 1/*λ*_*foot*_, with *λ*_*foot*_ = 36.31 µm. The measured ratio *λ*_*AP*_/*λ*_*foot*_ = 1.156 is within 0.8% of the predicted 1.165.

**Figure 4c** shows the corresponding axial current trace of the reference compartment of panel b and compares this with the axial current calculated from Equation (28) as *i*_*a*_ = *λ*_AP_*i*_*c*_, and the waveform from *i*_*c*_ = *c*_*m*_*dV*/*dt*. The superposition of the two curves confirms the relationship holds in this simulation.

**Figure 4d** demonstrates the spatial origin of the axial current at TCT1 in the reference compartment of the panel b fibre. To test Equation (31) we evaluated the membrane-current distribution at TCT1 during uniform propagation. Current sources behind the observation point were weighted by 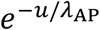 and integrated over the normalised distance *u*/*λ*_AP_. This reconstructed a local dV/dt of 940.844 mV ms^-1^ to within 0.08% of the value measured directly in the model 940.105 mV ms^-1^; the independent axial-current estimate *i*_*a*_/(*λ*_AP_*c*_*m*_) was 942.886 mV ms^-1^. Thus, for this uniformly propagating waveform, *λ*_AP_ quantifies the upstream distance over which distributed membrane current sources determine the local rate of depolarisation.

Figure 4 thus provides a numerical confirmation, in two model fibres with very different passive and active properties, of the relationships derived in Sections 2.2-2.5.

#### 2.7 The range over which *λ*_*AP*_ describes the AP upstroke

Equation (33) gives the local length 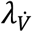 of the 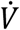 field at every point of the waveform, and Section 2.5 gives its limiting value in the ohmic foot. Together these conditionally bound the departure of the 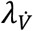 from *λ*_*AP*_ across the whole leading edge.

In an ohmic foot, 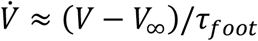 and *i*_*m*_ ≈ (*V* − *V*_∞_)/*r*_*m*_, so that the current ratio in Equation (33) takes the constant value:

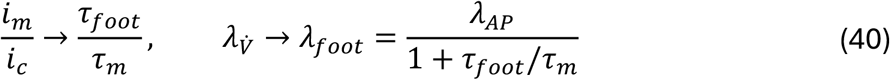

Between the fitted foot and TCT1, *i*_*m*_/*i*_*c*_ is expected to fall from this value to zero. If this condition holds, then with *ϵ* = *τ*_*foot*_/*τ*_*m*_, the fractional shortening of the local length relative to *λ*_*AP*_ over that interval is bounded, from Equations (33) and (40):

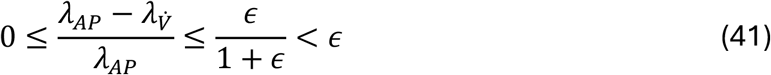

This quantifies the fractional error in using *λ*_*AP*_ as the local length scale during the leading edge of the AP, subject to the noted condition. For the baseline Hodgkin–Huxley fibre in Figures 1 and 2, *ϵ* = 0.0078 and the local *dV*/*dt* length therefore lies within 0.77% of *λ*_*AP*_ throughout the entire leading edge, including the foot. For the Markov fibre in Figures 3 and 4, *ϵ* = 0.165 and the corresponding bound is 14.2%. In a fibre of high passive resistance *λ*_*AP*_ may thus describe the whole leading edge to TCT1 with adequate accuracy; where resting leak is larger the discrepancy between the length scales is bounded and quantifiable.

Beyond TCT1 no analogous bound exists. Inward ionic current lengthens the local scale, and since 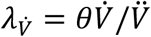 it diverges as 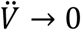 at peak *dV*/*dt*. However, it is possible to quantify the interval over which *λ*_*AP*_ remains an adequate descriptor of 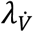. We define the normalized post-TCT extent 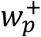 as the contiguous interval following TCT1 over which the local length remains within a fractional tolerance *p*:

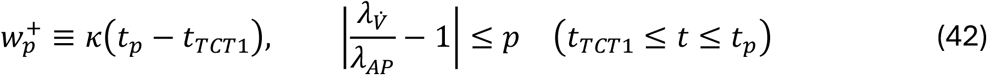

where *t*_*p*_ is the end of that interval. Because one unit of *κt* corresponds to one *λ*_*AP*_ in the travelling wave profile, the corresponding distance beyond TCT1 is 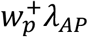.

The two measures, *ϵ* and 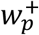, are formally independent. Toward the foot, spatial variation is bounded by *τ*_*foot*_/*τ*_*m*_, whereas towards the AP peak, it is determined by how rapidly inward current grows beyond the TCT. Both measures are therefore needed to state how much of the upstroke *λ*_*AP*_ describes. However, across the three fibres examined here they order consistently, from the Hodgkin–Huxley fibre (*ϵ* = 0.008, 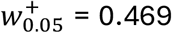) through a reduced-excitability Hodgkin–Huxley fibre (*ϵ* = 0.022, 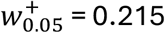; Section 4.4) to the Markov fibre (*ϵ* = 0.165, 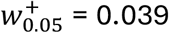), since both measures reflect the size of the ionic current relative to the capacitive current on either side of TCT1.

Sections 2.1 to 2.7 therefore establish *λ*_*AP*_ and *τ*_*AP*_ as exact constants of the uniform travelling wave, give their physical meaning, and relate them to routinely-measured waveform constants. We next describe the relationships between the AP constants and the classical passive constants.

### 3. Relationships between the DC, AC and AP constants

The constants developed in Section 2 describe a propagating AP waveform sustained by active ionic currents. The classical cable constants instead describe the response of a resting membrane to sustained or sinusoidal signals. We show in this section that they are exactly related to the AP constants. This demonstrates the two distinct contributions to conduction of active processes, quantified by *κ*, and of the passive cable that converts a waveform into distance and speed, quantified by diffusivity *D* = 1/(*r*_*a*_*c*_*m*_).

#### 3.1 The DC length and time constants

For a passive uniform cable near rest, a spatially uniform membrane perturbation charges exponentially with *τ*_*m*_, whereas the constant voltage produced by a maintained local current decays exponentially away from its source with *λ*_*DC*_ [1]. Combining equations (15) and (26) demonstrates that these constants share the same electrical diffusivity as the AP constants:

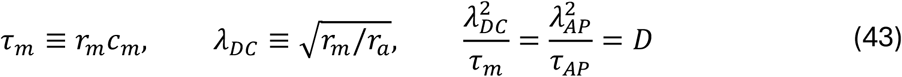

The active and passive pairs therefore relate as:

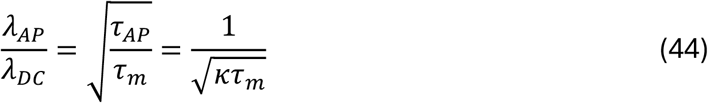

*λ*_*DC*_ and *λ*_*AP*_ are therefore diffusion lengths, 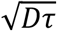, in the same physical cable, but evaluated over different timescales. Here the electrical diffusivity *D* = 1/(*r*_*a*_*c*_*m*_) characterises the electrical spread of charge, not the molecular diffusion of the ions themselves which is some four to five orders of magnitude slower in the baseline model fibre.

This reciprocal ratio also expresses conduction velocity in passive length constant units:

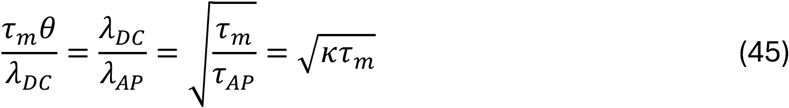

This dimensionless velocity has two geometric interpretations: how many AP lengths fit within one passive DC length, or how far a propagating AP waveform travels during one passive membrane time constant, in units of *λ*_DC_. It quantifies the separation between passive charging and the developing wavefront and thus has parallels with Rushton’s dimensionless quantity *S*_*R*_ = *α*_*R*_*θ*/*L*_*R*_ [1,12], although *α*_*R*_ and *L*_*R*_ are constants he defined that have parallels with *τ*_*m*_ and *λ*_*DC*_ but are not directly equivalent.

#### 3.2 The AC length constant

For angular frequency *ω*, the complex propagation constant of a uniform passive cable satisfies 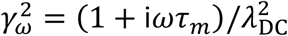, from a sinusoidal solution of Equation (13) [1,15]. The distance over which sinusoidal amplitude falls e-fold is given by the AC length constant, *λ*_AC_(*ω*) = 1/Re(*γ*_*ω*_). The imaginary part of the propagation constant quantifies the spatial phase lag. The exact comparison with the AP length follows from Equation (44) as:

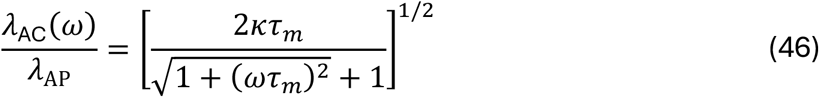

If an angular frequency is chosen such that *ω* = *κ*, the ratio between *λ*_AC_ and *λ*_AP_ approaches 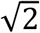 when *κτ*_*m*_ ≫ 1. This comparison explains the similarity between the scaling of AP and AC length constants at high-frequency passive cable scales. However, it should not be taken to mean that the AP can be described by a single sinusoidal frequency.

### 4. The determinants of *κ*, the AP constants and conduction reserve

The preceding sections separated the active waveform rate *κ* from the cable diffusivity *D* = 1/(*r*_*a*_*c*_*m*_). This allows biological and physical perturbations to be quantified separately by their influence on wavefront kinetics through *κ*, and by their influence on the passive properties that translate *κ* into wavefront distance and conduction velocity. The separation is one of mechanism rather than of biological independence: a change in *r*_*a*_ alters *D* while leaving *κ* nearly unchanged over the safely propagating range, whereas a change in *c*_*m*_ alters *D* directly and is also a determinant of *κ*.

#### 4.1 Na_*V*_ availability and membrane capacitance

At TCT1, *κ* = *d*ln(*dV*/*dt*)/*dt*, so *κ* measures the fractional rate at which the local rate of depolarisation is increasing. Maximum Na_*V*_ permeability and *c*_*m*_ were therefore varied separately to test how their balance determines *κ* (Figure 5a–c). We define the normalised ratio:

**Figure 5.**
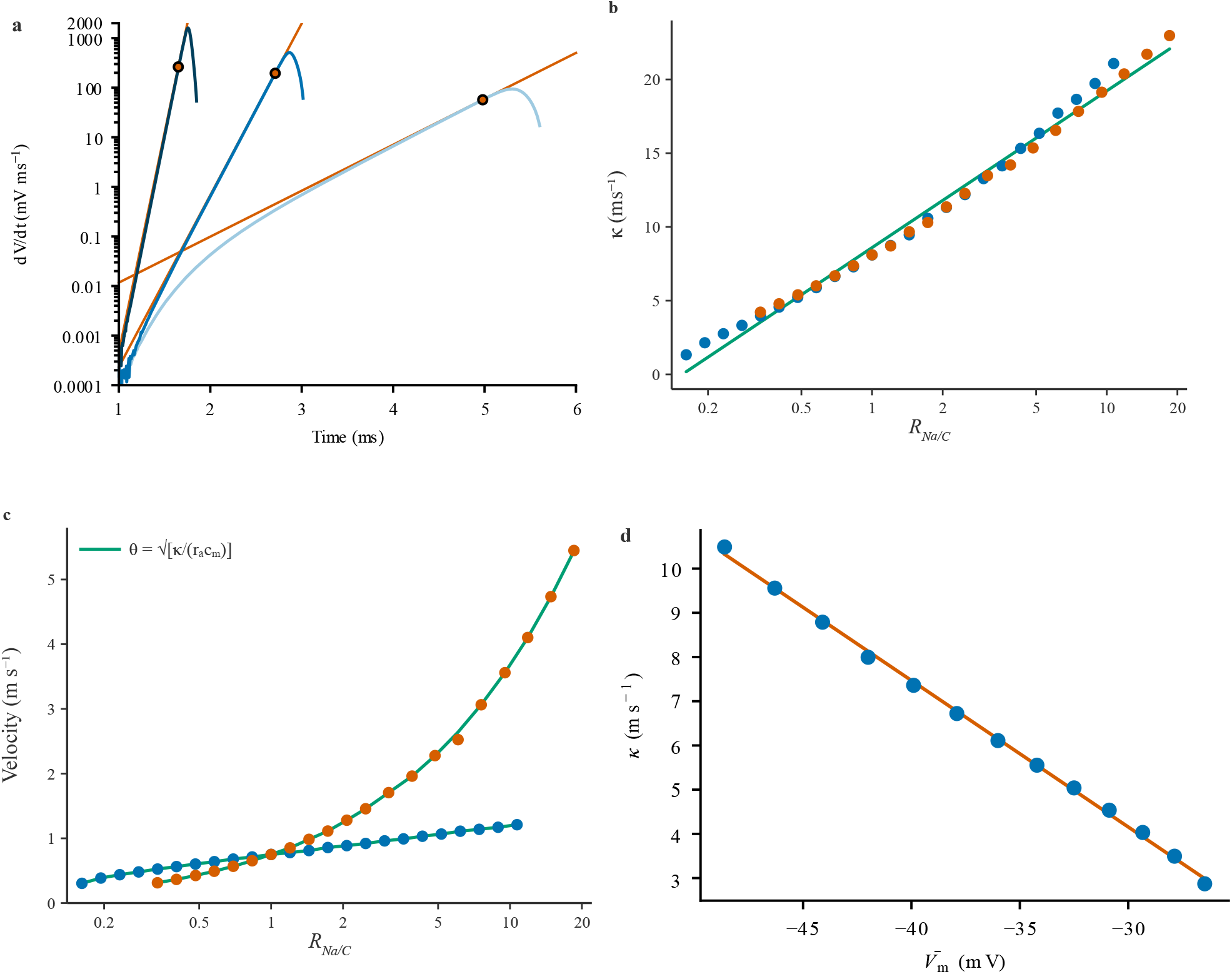
Dependence of κ and conduction velocity on Na_V_ and membrane parameters. **(a)** dV/dt against time for simulations with 10.5×, 1× and 0.23× baseline maximum Na_V_ permeability. Circles mark TCT1; orange lines show exponential reference functions with logarithmic growth rates κ = 20.9, 8.05 and 2.74 ms^−1^, respectively. Despite the marked curvature of the low-permeability trajectory, κ still predicts conduction velocity closely. **(b)** κ against the dimensionless ratio R_(Na/C_) for simulations varying P_NaV_(_max_) (blue) or c_m_ (orange). The green line is the pooled regression given in the text. **(c)** Measured conduction velocity for the same simulations (points) and the value calculated from 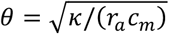 (green curves). **(d)** κ against the m-gate activation midpoint 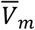.

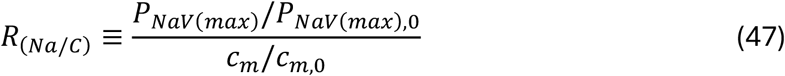

Here the subscript 0 denotes baseline model parameter values. Across the full range of propagating conditions in which either *P*_*NaV*(*max*)_ or *c*_*m*_ was varied while the other was held at baseline, *κ* followed the near log-linear empirical relation

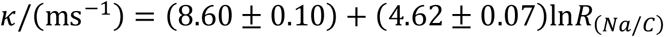

The fit gave *R*^2^ = 0.989, over the tested range 0.162 ≤ *R*_(*Na*/*C*)_ ≤ 18.5. Fits to the individual perturbation series gave similar slopes (4.60 ± 0.12 and 4.69 ± 0.09 ms^−1^ for varied Na_*V*_ permeability and *c*_*m*_ respectively). This fit is descriptive rather than mechanistic: its residuals are systematically curved, and at *R*_(Na/C)_ = 1 it lies approximately 7% above the baseline value of κ = 8.045 ms^−1^. Simulations outside the plotted range either failed to propagate or lacked a stable resting potential. This empirical relationship describes the tested perturbations; it does not establish a universal dependence on their ratio.

Figure 5c shows that despite similar influences on *κ*, proportional changes in *c*_*m*_ have a steeper influence on conduction velocity than *P*_*NaV*(*max*)_. This difference arises because increased Na_V_ permeability raises *κ* without changing *D*, thereby accelerating conduction while shortening 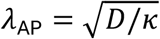. In contrast, reduced *c*_*m*_ increases both *D* and *κ*, thereby accelerating conduction but with much less influence on *λ*_AP_. Agreement between the measured velocities and 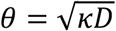 supports this relationship across the tested conditions.

#### 4.2 Voltage dependence of Na_*V*_ recruitment

The equivalent phase-plane expression 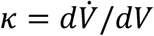 at TCT1 gives a complementary biological interpretation: *κ* measures how rapidly the rate of depolarisation increases with further depolarisation. Shifting the Na_*V*_ activation midpoint to more negative voltages recruits inward current earlier in the upstroke. In the tested simulations, *κ* decreased approximately linearly as the m-gate activation midpoint 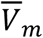 was shifted to more positive voltages (Figure 5d; slope −0.33 ms^−1^ mV^−1^, *R*^2^ = 0.999), with modest departure from linearity near the limits that permitted propagation.

#### 4.3 Axial and resting-membrane resistances

Axial resistance, *r*_*a*_, and resting membrane resistance, *r*_*m*_, influence conduction by different mechanisms. For a uniform continuum cable with unchanged membrane properties, changing *r*_*a*_ rescales distance and velocity without changing the steady travelling waveform in time. Increasing *r*_*a*_ therefore reduces *λ*_AP_ and velocity while leaving *κ* unchanged. Increasing diffusivity at fixed *κ* has the converse effect: both velocity and the AP length increase.

By contrast, *r*_*m*_ does not determine *D*, but influences *τ*_*m*_, *λ*_DC_ and the division of foot current between capacitive charging and ionic current. Sufficiently increased resting conductance can also alter the active waveform and prevent propagation.

Figure 6a and 6b illustrate these differences. Neither decreasing *r*_*m*_, by increasing resting chloride permeability, nor increasing *r*_*a*_ meaningfully influenced the phase-plane orbit or *κ* for parameter sets representing substantial propagation reserve. Instead, as predicted, increased axial resistance decreased the conversion of *κ* to velocity, and lower membrane resistance primarily increased foot current through the resting conductance. As *r*_*m*_ values neared the limits that permit successful propagation, phase-plane orbits contracted and TCT1 moved towards the point of maximum *dV*/*dt*. Here, the phase-plane slope tends to zero, such that *κ* and conduction safety fell until distal propagation failure occurred. Although no particular value of *κ* can be identified with conduction failure, its reduction reports the erosion of propagation safety reserve.

**Figure 6.**
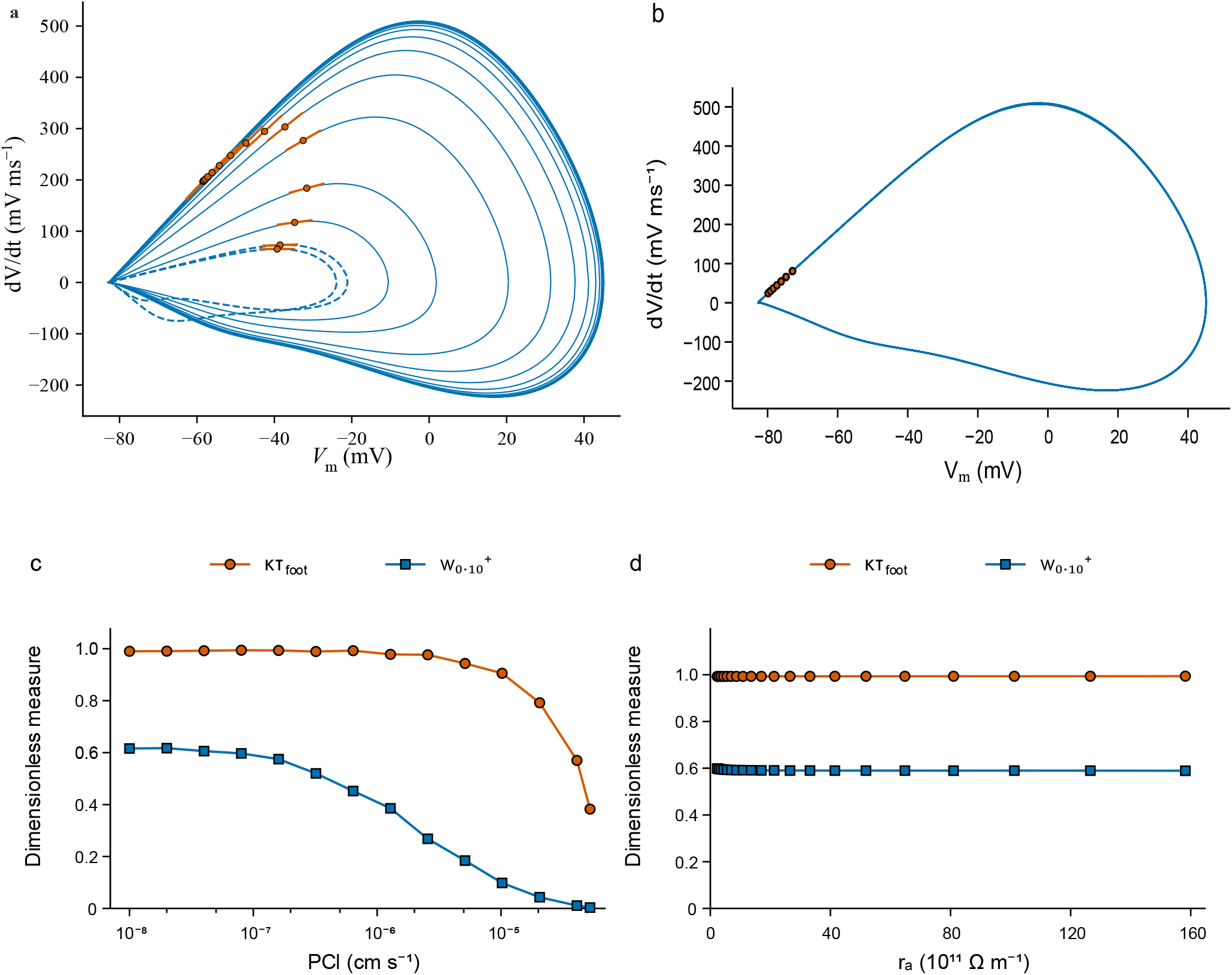
TCT position, κ and conduction reserve under passive loading. Panels **(a)** and **(b)** show phase-plane trajectories of dV/dt against V for model fibres in which: **(a)** resting membrane resistance r_m_ was reduced by increasing chloride permeability. The outer 13 phase plots show the effect of repeatedly doubling the membrane Cl^-^ permeability from the default value r_m_ = 2.53 x 10^5^ Ω m (outermost plot, leftmost TCT) to 49,037 Ω m. The innermost three plots show smaller decreases in r_m_, to 38499, 36747 and 36738 Ω m respectively. Dashed trajectories denote simulations in which the AP attenuated with distance and did not reach the end of the model fibre. Circles mark TCT1 and orange lines show local phase-plane tangents, of slope κ. **(b)** axial resistance r_a_ was changed over a range from r_a_ = 2.28 × 10^11^ to 1.582 × 10^13^ Ω m^−1^. Markers denote TCT1. Waveform shape and κ changed little across conditions with stable propagation. As the propagation boundary approached, the orbits contracted and TCT1 moved towards the maximum of dV/dt, reducing κ. **(c)** and **(d)** κτ_foot_ (the capacitive fraction of foot current) and 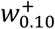 (the normalised distance beyond TCT1 over which 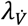 remains within 10% of λ_AP_) for the uniformly conducting simulations in (a) and (b), plotted against the corresponding parameter.

#### 4.4 The influence of membrane and axial resistance on the range over which *λ*_AP_ describes the wavefront

The two measures developed in Sections 2.5 and 2.7 (Equations (38) and (42)) were applied to the uniformly conducting simulations of Figure 6: *κτ*_*foot*_, the fraction of converging axial current that charges the membrane capacitance in the exponential foot, and 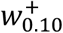, the normalised distance beyond TCT1 over which 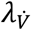 remains within 10% of *λ*_*AP*_. As resting chloride permeability increased (Figure 6a, c), *κτ*_*foot*_ remained close to unity and then fell steeply from 0.99 to 0.38, while 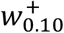 declined more steadily from 0.62 to 0.004. By contrast, as *r*_*a*_ changed (Figure 6b, d), *κτ*_*foot*_ remained close to unity and 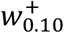 remained in the range 0.589–0.599, consistent with axial resistance acting primarily through cable conversion. Both measures are computable from a single voltage trace once κ is known; together they quantify foot current partition and the post-TCT1 interval over which *λ*_*AP*_ approximates the local length scale.

### 5. Recovery of *κ* from the voltage waveform and *τ*_*m*_

κ is defined at a TCT, and locating a TCT formally requires knowledge of *i*_*m*_. Equation (37) provides a mathematically exact alternative requiring only the foot exponent and an independent estimate of *τ*_*m*_. We tested it against varied model waveforms for which the true value of *κ* is known.

In an ohmic exponential foot, *V* − *V*_∞_ ∝ *e*^*αt*^ with *α* = 1/*τ*_*foot*_. This exponent may be obtained from the slope of 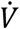 against *V* − *V*_∞_ with the fit constrained through the resting point, or from the slope of 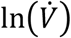 against time. Applied to the AP of Figure 1, for which κ = 8.045 ms^−1^ had been determined directly at the TCT from the simulated current waveform, a chord fit to the foot of the voltage trace alone gave α = 8.105 ms^−1^; Equation (37) with the model *τ*_*m*_ = 15.87 ms returned κ = 8.042 ms^−1^, recovering the known value to within 0.04%. For the Markov model of Figure 3, *α* = 16.36 ms^-1^ and *τ*_*m*_ = 0.371 ms, giving *κ* = 14.044 ms^-1^, within 0.7% of *κ* = 14.143 ms^−1^ measured at TCT1.

Equation (37) is robust to the value of *τ*_*m*_, potentially allowing an estimated value to be used in some circumstances. Logarithmic differentiation of Equation (37), holding the measured *α* fixed, shows that the relative sensitivity of *κ* to the assumed *τ*_*m*_ is ∂ln*κ*/ ∂ln*τ*_*m*_ = 1/(1 + *ατ*_*m*_). This is small whenever the foot is fast relative to passive charging.

Recovery of *κ* therefore requires only a voltage trace, a value for *τ*_*m*_, and the assumption of an approximately ohmic foot across the fitted interval. It does not require that a phase plot is linear near the TCT itself. The application of this procedure to experimental recordings, and the further quantities that *κ* then makes accessible, are the subject of a future study.

## Discussion

For more than a century, cable theory has provided the mathematical language for understanding voltage spread in neurons and other biological cables; Rall offers a brief, insightful history of its development [4]. A particular strength of the theory is that it expresses passive membrane properties through two intuitive constants, *λ*_*DC*_ and *τ*_*m*_, quantities that have been measured and reasoned with since the 1940s [2]. The propagating action potential has proved less tractable: from the first travelling-wave treatments [12] through the numerical solution of the propagated impulse [3] to the analytical and reduced models that followed [1,13,14], conduction velocity has been recovered by numerical search or by approximation, and it has not been possible to describe the active wavefront with characteristic lengths or times analogous to the passive constants. Here we show that the AP wavefront can be described by exact length and time constants, that each has a precise physical meaning, and each can in principle be recovered from voltage traces.

The derivation exploits specific instants during propagation that we term transmembrane current transition (TCT) points. In a uniformly propagating AP, the net transmembrane ionic current must change sign at least twice. At these points, the cable equation reveals a rate coefficient *κ* = *θ*^2^*r*_*a*_*c*_*m*_ that may be read from the waveform as *κ* = (*d*^2^*V*/*dt*^2^)/(*dV*/*dt*) at a TCT, or equivalently as the slope of the dV/dt vs V phase plane at that point. Although defined at an instant, *κ* is a single-valued constant of a uniform waveform.

*κ* describes the instantaneous logarithmic rate of |*dV*/*dt*| at a TCT, so we define its reciprocal as the action potential time constant, *τ*_*AP*_ = 1/*κ*. The travelling-wave relation ∂/ ∂*t* = −*θ* ∂/ ∂*x* converts this rate into the action potential length constant 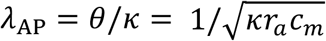, giving the exact expression *θ* = *λ*_AP_/*τ*_*AP*_. A uniformly propagating action potential travels a distance of one *λ*_AP_ in a time of one *τ*_*AP*_.

Because the amplitude of a propagating AP does not decay with distance, *λ*_AP_ does not describe attenuation as the steady state length constant *λ*_*DC*_ does. Instead, it may be considered to exactly describe three aspects of the propagating waveform.

1. *λ*_AP_ = *i*_*a*_/*i*_*c*_ throughout the waveform, wherever *dV*/*dt* ≠ 0. This is the active counterpart of *λ*_*DC*_ = *i*_*a*_/*i*_*m*_ over a steady-state exponential profile.
2. *λ*_AP_ is the range over which transmembrane ionic current sets local *dV*/*dt*, because local capacitive current at any point is equal and opposite to an exponentially weighted average of the ionic current on the excited membrane behind it, with an *e*-folding distance of *λ*_*AP*_. This weighting is exponential even where the waveform is not, giving *λ*_*AP*_ an exact spatial meaning at every point of the wave. This is the active counterpart of *λ*_*DC*_ describing the *e*-folding length of a symmetric kernel.
3. *λ*_AP_ is the exact local length scale of the *dV*/*dt* field at a TCT, and hence of the capacitive and axial current fields that are proportional to it.

In all parts of the waveform other than the TCTs this third equality becomes an approximation, the accuracy of which may be quantified. Toward the foot from TCT1, the local length of the dV/dt field tends towards *λ*_foot_. If 0 ≤ *i*_*m*_/*i*_*c*_ ≤ *τ*_foot_/*τ*_*m*_ throughout this interval, its fractional shortening relative to *λ*_AP_ is at most *τ*_foot_/(*τ*_*m*_ + *τ*_foot_), and is therefore less than *τ*_foot_/*τ*_*m*_. Beyond TCT1 the local length becomes unbounded, since inward current lengthens the local scale as *dV*/*dt* approaches its maximum. Instead, we describe the normalised extent *w*_p_^+^ to quantify how far past TCT1 *λ*_AP_ describes the local spatial scale to a chosen tolerance. *λ*_AP_ is therefore a quantity with an exact local meaning and a stated, measurable range of applicability.

*λ*_AP_ and *τ*_*AP*_ have exact relationships with the classical cable constants that help clarify their physical meaning. *λ*_*DC*_ and *λ*_AP_ are both diffusion lengths 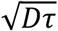 in the same cable but evaluated over different times, so that 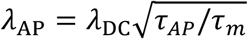. Electrical diffusivity *D* = 1/(*r*_*a*_*c*_*m*_) describes electrical spread along the cable; the two length constants differ according to their timescale ratio, with *λ*_AP_ shorter than *λ*_*DC*_ when *κτ*_*m*_ > 1. We show that the AC length constant, *λ*_*AC*_ exceeds *λ*_AP_ by a factor approaching 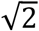 when the imposed angular frequency is set equal to *κ* and *κτ*_*m*_ ≫ 1. The foot exponent 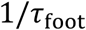 [1] is similarly related to *κ* without equalling it. Under the classical ohmic-foot assumption *τ*_AP_ = *τ*_foot_(1 + *τ*_foot_/*τ*_m_) so the foot exponent always exceeds *κ* by the fraction *τ*_foot_/*τ*_m_. Where *τ*_m_ ≫ *τ*_foot_ the two may differ negligibly.

The separation of active and passive properties suggested by the expression of *λ*_*AP*_ in terms of *κ*, describing the active waveform, and the diffusivity, *D*, describing the cable properties, naturally divides the determinants of conduction velocity into two groups that can be examined separately. We show that Na_V_ properties act on *κ* alone, axial resistance acts on *D* alone, and membrane capacitance is the only parameter that directly acts on both. Thus, raising Na_V_ permeability accelerates conduction while shortening *λ*_*AP*_, whereas lowering *c*_*m*_ accelerates conduction with much less effect on *λ*_*AP*_, since it raises *κ* and *D* together. In principle this allows an observed change in velocity to be attributed to membrane excitability or to axial coupling rather than left as a composite of the two, and it offers quantities with which to examine, for example, developmental changes in axon diameter and conduction speed, such as those demonstrated in myelinated CNS axons [23] or the activity-dependent changes in conduction velocity reported in small-fibre neuropathy [24].

The same quantities may be useful in reporting approaching conduction failure. When resting membrane resistance was reduced towards values at which distal propagation failed, the phase-plane orbits contracted, TCT1 moved towards the point of maximum dV/dt, and κ decreased. Under progressively increased resting leak, *w*_p_^+^ fell substantially while *κτ*_foot_ was still close to unity, so the two appear to degrade in a defined order as the boundary is approached. Neither identifies a threshold value of *κ* at which conduction fails, but both make the erosion of conduction reserve visible in the waveform before failure occurs.

Application of the concepts developed here requires some method to determine *κ* experimentally, yet locating a TCT formally requires knowledge of the *i*_m_ waveform. The relationship of *κ* to the foot time constant provides a solution. Thus, *κ* = *α*^2^*τ*_*m*_/(1 + *ατ*_*m*_) requires determination of only the foot exponent *α* = 1/*τ*_foot_ and an estimate of *τ*_*m*_, both of which are routine measurements. It does not require TCT1 to sit on a linear part of the phase plane plot. The relative sensitivity of *κ* to the assumed *τ*_*m*_ is 1/(1 + *ατ*_*m*_).

The key findings were tested in model fibres using both Hodgkin-Huxley and Markov ion channel formalisms and across the full range of channel densities that were compatible with AP propagation. Waveform-derived values of *κ* predicted conduction velocity over more than a fivefold range of velocities. The set of relationships between *κ*, the action potential constants and conduction velocity is not dependent on model formalisms.

This study has not attempted to study these relationships in an experimental preparation. However, the measurements this approach requires have been made for decades. Phase plots of the kind shown in Figures 2, 3, and 6 have a long history in membrane theory [25] and in experimental studies of skeletal muscle [26] and cardiac muscle [27], and remain in routine use in, for example, feline cortical neurons [28] and murine hippocampal interneurons [29]. Given a suitable foot and an independent estimate of the relevant *τ*_*m*_, κ may in principle be estimated retrospectively from a voltage record. Multielectrode recordings of the kind that would allow *λ*_*AP*_ and/or velocity to be measured directly alongside AP waveforms are already commonplace in larger fibres such as skeletal muscle [19,30]. Indeed, because *κ* is a logarithmic rate, it is unchanged by constant scaling or offset of the voltage signal, potentially allowing its recovery from extracellular recordings that preserve the relevant upstroke dynamics. Such intracellular AP upstroke features may be derived from extracellular potentials in some systems [31].

A further limitation of the present study is that the derivations assume a uniform, unterminated, unmyelinated cable carrying a steadily propagating action potential, and real axons, dendrites and muscle fibres may depart considerably from that idealisation. Tapering, branching, myelination and the skeletal muscle t-system all require position-dependent constants in place of the single global constant appropriate to a uniform cable [15,32,33]. Because *κ* is the local real exponent of the propagating wave, the natural extension is to evaluate the geometry-specific spatial problem at that exponent, but a fully tested quantitative theory for each geometry must remain beyond the present scope. Finally, recovery of *κ* from *τ*_foot_ assumes an approximately ohmic foot across the fitted interval, an assumption that weakens as resting leak becomes a larger fraction of total conductance and which *κτ*_foot_ itself quantifies. Gating dynamics can contribute frequency-dependent membrane admittance [34], such that the relevant *τ*_*m*_ might not equal a time constant measured under different conditions.

In summary, by identifying and exploiting transmembrane current transition points, we have derived exact length and time constants for the propagating action potential and established their precise relationships with conduction velocity, with the classical DC, AC and foot constants, and with the local length scales of the AP wavefront. The rate coefficient *κ* is compact, exactly defined, independent of the gating formalism, and in principle measurable from an ordinary voltage trace. Almost ninety years after Rushton asked what determines the speed of the propagated disturbance [12], the uniformly propagating action potential can be described in the same intuitive language of length and time constants that has served the passive membrane so well.

## Additional Information

JA Fraser conceived of the project, developed the model employed, derived the mathematical proofs, conducted and interpreted simulations and drafted the manuscript.

E Lopez-Belmonte Deza conducted and interpreted simulations and critically revised the manuscript.

Both authors approved the final version of the manuscript; agree to be accountable for all aspects of the work in ensuring that questions related to the accuracy or integrity of any part of the work are appropriately investigated and resolved; and qualify for authorship. All those who qualify for authorship are listed.

The work was not funded by a research grant.

The authors have no competing interests.

