## Supplementary Methods for "The length constant, time constant and velocity of a propagating action potential"

### Supplementary Methods for the Markov Model

The Markov simulations used the charge-difference model described in the Methods for the calculation of membrane potential, Nernst–Planck axial electrodiffusion, $Na^+/K^+$-ATPase, ionic concentration and volume equations. The voltage-gated membrane channel formalisms and background ion conductances were altered as described below, giving different steady state initial ionic composition. Figure-specific equilibration, cable and measurement protocols differed as detailed below.

#### Voltage-gated conductances

NaV membrane current was represented by the deterministic six-state human NaV1.6 scheme of Balbi et al., comprising two closed (C1, C2), two open (O1, O2) and two inactivated (I1, I2) states connected by 12 voltage-dependent transitions. The published topology and transition-rate parameters were reimplemented in the charge-difference solver without fitting or alteration. State fractions were advanced by forward Euler at each adaptive time step; negative round-off values were clipped and all six fractions renormalised to conserve total occupancy. The implementation applied a $Q_{10}$ of 3 to transition rates; simulations were performed at its 293.15 K reference temperature, so this factor was unity.

To preserve the conductance formulation of the source model, sodium current was calculated from the summed open probability O1 + O2, the specified maximum conductance and the instantaneous sodium equilibrium potential; in contrast with the HH formalisms in the main text, this was not converted to a Goldman permeability. Repolarising current used the Dodge–Cooley delayed-rectifier formulation that provided the potassium-conductance context for Balbi's hybrid axon: an ohmic potassium conductance gated by $n^4$, with the rate argument V − Vrest − Vshift, where Vrest = −80 mV and Vshift = −10 mV. Kinetic equations were transcribed from the corrected dc.mod implementation archived as ModelDB model 3805 (repository commit 6ee0e303f67008bb9c4569b315b7a4e67170037e) and were not refitted. The permeability-based Hodgkin–Huxley membrane $Na^+$ and $K^+$ currents were disabled in these runs; the electrodiffusive axial and charge-difference calculations were unchanged.

#### Parameter sets

The conductance densities used in Balbi's NaV1.6/Dodge–Cooley hybrid simulation (1.20 S $cm^{-2}$ NaV1.6 and 0.070 S $cm^{-2}$ K channel) defined the reference scale. The NaV–capacitance series retained this K channel density and used 0.90×, 1.00× or 1.10× the

NaV density. The high-resolution phase-plane and spatial-wavefront reference used 1.15× NaV and 0.50× K to provide a distinctly curved upstroke while retaining stable, uniform propagation. No transition rate was changed. Simulations used the Thio et al. unmyelinated-fibre ionic milieu: extracellular [$Na^+$], [$K^+$] and [$Cl^-$] were 154, 5.4 and 159.4 mM, and initial intracellular values equilibrated at 8.9, 145 and 6.715 mM, respectively, for the baseline parameter set. Ionic diffusivities were scaled uniformly by 0.851146 to target an axial resistivity of 100 Ω cm. The membrane leak currents were ohmic. The reference stimulus permeability amplitude was 3.84 × $10^{-4}$ cm $s^{-1}$; in the NaV–capacitance series it was reduced to 0.25–1.00× this value according to capacitance to avoid stimulus-induced instability.

**Supplementary Table S1.** Parameters specific to the final Markov figure datasets.

| Parameter | High-resolution reference | NaV–capacitance series |
|---|---|---|
| **Temperature** | 293.15 K (20 °C) | 293.15 K (20 °C) |
| **Fibre diameter** | 1.0 μm | 1.0 μm |
| **Membrane capacitance** | 1.000 μF $cm^{-2}$ | 0.119–1.948 μF $cm^{-2}$ |
| **Maximum NaV1.6 conductance** | 1.38 S $cm^{-2}$ (1.15×) | 1.08, 1.20 or 1.32 S $cm^{-2}$ (0.90×, 1.00× or 1.10×) |
| **Maximum Dodge–Cooley K conductance** | 0.035 S $cm^{-2}$ (0.50×) | 0.070 S $cm^{-2}$ (1.00×) |
| **Na leak conductance** | 4.20484 × $10^{-5}$ S $cm^{-2}$ | 4.0554–4.3541 × $10^{-5}$ S $cm^{-2}$ |
| **K leak conductance** | 1.14300 × $10^{-3}$ S $cm^{-2}$ | 1.14146–1.15806 × $10^{-3}$ S $cm^{-2}$ |
| **Cl leak conductance** | 1.000 × $10^{-3}$ S $cm^{-2}$ | 1.000 × $10^{-3}$ S $cm^{-2}$ |
| **Initial [Na]/[K]/[Cl], outside; inside** | 154/5.4/159.4; 8.9/145/6.715 mM | 154/5.4/159.4; 8.9/145/6.715 mM |
| **Cable discretisation** | 600 × 4.165 μm | 80 × 8.33 μm |

#### Resting state and restart protocol

The non-negative Na and K leak conductances were first obtained for the published-density condition by balancing the pump and resting voltage-gated fluxes at −80 mV; Cl conductance was fixed at 1.0 × $10^{-3}$ S $cm^{-2}$, at which the initial chloride equilibrium potential was approximately −80 mV. The high-resolution reference retained these validated leak conductances after its active Na and K densities were changed, and the full model was allowed to find its own equilibrium rather than forcing the membrane potential to remain at −80 mV. It consequently rested at −78.5098 mV. In the NaV–capacitance series, intracellular impermeant charge was adjusted so that the common starting ionic composition corresponded to −80 mV, and the passive Na and K conductances were rebalanced independently for each active-channel density before equilibration; all accepted values were non-negative. The reference fibre's frozen-state differential membrane resistance, measured from the centred total-current response to ±0.1 mV with concentrations and channel states held fixed, was 370.61 Ω $cm^2$; with 1.0 μF $cm^{-2}$ capacitance this gave $\tau_m$ = 0.37061 ms. Its effective axial resistivity was 99.674 Ω cm (target 100 Ω cm).

Each distinct parameter set was equilibrated independently in one unstimulated compartment using successive 250 ms restart segments. At least 2 s was simulated, with a 20 s ceiling for the NaV–capacitance series. Equilibrium required two consecutive segments in which the voltage changed by <10 μV, each mobile-ion concentration by <1 μM, every Markov occupancy by $<10^{-6}$, the state sum differed from unity by $<10^{-9}$ and net membrane current was <0.01 μA $cm^{-2}$. The complete accepted state was then copied identically to every cable compartment and time was reset. Before stimulation, each cable was run for 20 ms and required <10 μV temporal voltage drift and <1 μV spatial variation. The high-resolution reference met the single-compartment criteria after 6.0 s.

### Cable simulations and numerical verification

For the high-resolution spatial analysis, the 1 μm-diameter cable contained 600 compartments of length 4.165 μm (total length 2.499 mm), with a maximum time step of 25 ns and a 1 μs saved waveform interval. Voltage and $\dot{V}$ were saved at every requested spatial site.

Before selecting the altered high-resolution reference, the published-density NaV1.6/Dodge–Cooley condition was tested in the same 2.499 mm physical cable at compartment lengths of 8.33, 4.165 and 2.0825 μm and maximum time steps of 50 and 25 ns. All six combinations retained uniform propagation, and every prespecified waveform and propagation metric differed by <2% between the two finest spatial or temporal resolutions. This check validated the numerical implementation but was not a separate convergence series for the subsequent 1.15× NaV/0.50× K reference. In that selected 4.165 μm reference cable, relative ranges across the central uniform region were 0.001% for velocity, <0.001% for peak amplitude and action-potential duration, 0.011% for maximum upstroke and 0.494% for $\kappa$ at TCT1. Independent differentiation of stored voltage agreed with the model-reported $\dot{V}$ with 0.005% median and 0.151% 95th-percentile relative error where $|\dot{V}| \geq 0.01$ V $s^{-1}$.

The NaV–capacitance comparison comprised 11 independently equilibrated 1 μm-diameter cables, each with 80 compartments of length 8.33 μm. The condition-dependent maximum time step was 5–50 ns. Membrane capacitance ranged from 0.119 to 1.948 μF $cm^{-2}$ and was crossed with the three NaV density levels specified above to span the plotted range without changing Markov kinetics. Waveforms were saved at compartments 20, 30, 38–42, 50 and 60 in the verification records, and the central site (compartment 40) was used for the phase-plane measurement after confirming agreement of local propagation intervals.

#### Extraction of transition and propagation quantities

TCT1 and TCT2 were the successive sign changes of total transmembrane ionic current from net outward to inward and from net inward to outward, respectively. For the high-resolution reference, $\kappa$ at TCT1 was evaluated from the derivatives of a quartic fit to $V(t)$ over ±30 µs. For the velocity series, $\kappa$ was recalculated from the raw saved waveform. $\dot{V}$ was differentiated from stored $V$ before calculating the ordinary-least-squares slope of $\dot{V}$ against $V$ over the five samples bracketing TCT1. Measured conduction velocity was the reciprocal of the slope obtained by regressing interpolated 0 mV arrival time against distance across compartments 38–42, and was accepted only when it agreed with estimates from remote recording sites. Calculated velocity used the relationship given in the main text with the measured $\kappa$, membrane capacitance and local axial resistance.

For the spatial-wavefront analysis, all fields were taken from a single saved instant at the reference-compartment TCT1. The local length of the $\dot{V}$ field was evaluated as $-\dot{V}/(\partial\dot{V}/\partial x)$, using a symmetric fourth-order five-point spatial derivative; a three-point centred derivative provided a discretisation check. The temporal foot exponent $\alpha = 1/\tau_{\text{foot}}$ was obtained by linear regression of $\ln(\dot{V})$ against time before TCT1, using samples for which $\dot{V}$ was 5–50% of its value at TCT1; the corresponding spatial foot slope was $\alpha/\theta$. Where the post-TCT width was reported, the local temporal slope $S_1$ was the five-point regression slope of $\dot{V}$ against $V$, and $w_p^+ = \kappa(t_p - t_{\text{TCT1}})$ was the first contiguous post-TCT1 interval satisfying $|\kappa/S_1 - 1| \le p$.
